# BASTA, a simple whole blood assay for measuring β-cell antigen specific CD4^+^ T-cell responses in type 1 diabetes

**DOI:** 10.1101/2025.01.08.631617

**Authors:** Matthew Lacorcia, Pushpak Bhattercharjee, Abby Foster, Melinda Y. Hardy, Jason Tye-Din, John Karas, John M Wentworth, Fergus Cameron, Stuart I. Mannering

## Abstract

Type 1 diabetes (T1D) is an autoimmune disease where T-cells mediate the destruction of the insulin-producing β-cells found within the Islets of Langerhans in the pancreas. Currently, autoantibodies to β-cell antigens are the only tests available to detect β-cell autoimmunity. T-cell responses to β-cell antigens, which are known to cause T1D, can only be measured in research settings due to the complexity of assays and the large blood volumes required. Here we describe the <u>β</u>-cell <u>a</u>ntigen <u>s</u>pecific <u>T</u>-cell <u>a</u>ssay – BASTA. BASTA is a simple whole blood assay that can detect human CD4^+^ T-cell responses to β-cell antigens by measuring antigen-stimulated IL-2 production. BASTA is both more sensitive and specific than the CFSE-based Proliferation Assay. We used BASTA to identify the regions of preproinsulin which stimulated T-cell responses specifically in blood from people with T1D. BASTA can be done with as little as 2-3 mL of blood. We found that effector memory CD4^+^ T cells are the primary producers of IL-2 in response to preproinsulin peptides. We then evaluated responses to individual and pooled preproinsulin peptides in a cross-sectional study of pediatric subjects: without T1D, without T1D but with a first-degree relative with T1D, or diagnosed with T1D. In contrast to other preproinsulin peptides, full-length C-peptide (PI_33-63_) showed excellent specificity for T1D (AUC = 0.86). We suggest that BASTA will be a useful tool for monitoring changes in β-cell specific CD4^+^ T-cell responses both in research and clinical settings.

**One Sentence Summary:** β-cell antigen specific CD4^+^ T-cell responses in children with T1D can be measured using a simple whole blood assay.

## INTRODUCTION

Type 1 diabetes (T1D) is a T-cell-mediated autoimmune disease arising from destruction of pancreatic insulin-producing β-cells [1, 2]. β-cell antigen-specific CD4^+^ and CD8^+^ T cells collaborate to mediate β-cell destruction [1]. However, CD4^+^ T-cells are thought to play a central role in the autoimmune pathogenesis because of the high risk for developing T1D that is associated with certain Human Leukocyte Antigen (HLA) class II alleles [3, 4]. A simple, sensitive and robust assay to measure the frequency and function of these β-cell antigen-specific CD4^+^ T cells has been long sought after by the T1D community [5]. However, despite much effort, there is currently no assay suitable for routine clinic-based monitoring of these rare autoantigen-specific cells.

Anti-β-cell autoimmunity is currently detected by measuring β-cell specific autoantibodies. These autoantibodies are believed to reflect an underlying pathogenic autoimmune T-cell response [4]. Currently, autoantibodies to four β-cell proteins are routinely measured: insulin, glutamic acid decarboxylase 65kDa (GAD-65), islet tyrosine phosphatase (IA-2) and zinc Transporter 8 (ZnT8) [6]. These are collectively referred to as ‘islet autoantibodies’. An individual’s risk of developing clinical T1D increases as more β-cell proteins are targeted by autoantibodies [4], with the appearance of multiple islet autoantibodies now designated as “Stage 1” T1D [7]. Paradoxically, while islet autoantibodies are a useful marker for identifying individuals at high risk of developing T1D, they are not believed to have a direct role in the pathogenesis of T1D [8, 9]. Furthermore, islet autoantibodies are poor predictors of when an individual will progress from normal glucose tolerance to dysglycemia (“Stage 2” T1D) and ultimately insulin dependent, clinical (or “Stage 3”) T1D. T-cell assays may better reflect the underlying immunopathogenesis of T1D. β-cell antigen specific are attractive targets for emerging disease-altering therapies for T1D [10], yet we remain largely blind to T-cell responses due to the logistical and technical challenges inherent in measuring β-cell antigen-specific T-cells in T1D.

The detection and analysis of T-cell responses associated with T1D in a clinical setting faces two major challenges. First, the low frequency of β-cell antigen-specific T cells in the peripheral blood, estimated to be in the order of 1:10^5^-10^6^ T cells [11]. The second challenge is that the β-cell antigens recognized by pathogenic CD4^+^ T cells in people with T1D are poorly characterized, although considerable progress has been made in this area over recent years [12–14].

Despite these challenges, four assay types are currently used to detect β-cell antigen-specific CD4^+^T cells in peripheral blood [11, 15, 16]. They are: (i) the CFSE-based proliferation assay (and related dye dilution assays) [17], (ii) ELISpot assays [18], (iii) tetramer staining [19] and (iv) activation marker upregulation [20]. The activation-induced marker (AIM) assay, which uses flow cytometry to detect the upregulation of cell surface activation markers, is the most recently developed [20]. Each of these assays has contributed to our understanding of T-cell responses in T1D, but face significant intrinsic obstacles as screening tools for widespread clinical use [16], including: extensive laboratory-based preparation, large volumes of blood, the need for highly skilled technicians, and in many cases customized reagents and/or sophisticated bioinformatic analysis. Logistically, many of these assays are problematic because freezing/thawing damages some cell populations, particularly antigen-experienced T cells [21], which compromises shipping and batch analysis of samples.

The analysis of antigen-specific CD4^+^ T-cell function requires knowledge of the appropriate antigens [5]. It is now clear that insulin and its precursor preproinsulin is a central antigen in the autoimmune pathogenesis of T1D [14, 22]. More recently, C-peptide from proinsulin has emerged as a major antigen in human T1D. It is recognized by both human islet-infiltrating [12] and peripheral blood CD4^+^ T cells [13, 23, 24], and elicits responses associated with disease progression [25]. In addition, responses to insulin B-chain [26] and A-chain have been well documented [27, 28] in the blood of people with T1D.

Peripheral blood is the only tissue that is routinely available to measure human immune responses. Whole blood assays have been used to detect T-cell responses to the vaccine antigens tetanus toxoid [29] and SARS-CoV-2 [30] by measuring cytokines in the plasma after a period of *in vitro* culture. In fact, stimulation of whole, heparinized blood, has become a standard diagnostic to detect responses to mycobacterial infection [31]. Despite their utility, these whole blood functional antigen-specific T-cell assays were not sufficiently sensitive to measure autoimmune T-cell responses. However, the recent development of very sensitive ELISA-like assays that use electrochemiluminescence (ECL)-based cytokine detection methods has facilitated the development of whole blood assays capable of measuring autoimmune CD4^+^ T-cell responses. A whole blood assay combined with very high-sensitive cytokine detection has recently been used to detect responses to gliadin peptides in celiac disease [32, 33]. Here we describe the development and validation of a simple whole blood assay capable of detecting β-cell antigen-specific CD4^+^ T-cell responses well suited to use in a clinical setting, which we call BASTA (<u>β</u>-cell <u>a</u>ntigen <u>s</u>pecific <u>T</u>-cell <u>a</u>ssay).

## RESULTS

### BASTA is superior to the CFSE-based Proliferation Assay

Previously we have used the CFSE-based Proliferation Assay to measure CD4^+^ T-cell responses to full-length C-peptide from proinsulin (PI_33-63_) [12, 13, 17]. To evaluate BASTA, we first compared CD4^+^ T-cell responses to full-length C-peptide detected using the CFSE-based Proliferation Assay to responses detected using BASTA from the same blood samples (Fig. 1A). In the CFSE-based Proliferation Assay, the number of cells that proliferate in response to an antigen over 7 days are counted by flow cytometry. Responses are expressed as a ratio (cell division index, or CDI) of the number of cells that have proliferated with and without antigen [17]. For this cohort, while the CDI values were increased in the T1D group (n = 17) compared to the non-T1D samples (n = 8), this was not statistically significant (Mann-Whitney, *P* = 0.92) (Fig. 1B). In contrast, using BASTA C-peptide specific responses were determined by measuring the concentration of IL-2 present in the plasma after 24-hour of autoantigen stimulation, in the high sensitivity Meso Scale Discovery (MSD) electrochemiluminescence-based immunoassay platform [33]. We chose to measure IL-2 because this cytokine has a very low basal concentration and consequently, gives the best signal-to-noise ratio compared to IFNγ, TNFα, IL-6, IL-10, and IL-13 (Supplementary Figure 1C). In addition, IL-2 was shown to be a useful readout for measuring CD4^+^ T-cell responses to gliadin peptides in individuals with celiac disease [32]. The positive control was the vaccine antigen tetanus toxoid (TT), which gave comparable results to a commercially available pool of viral peptides (CEFX) (Fig. S1B-C). Antigen-stimulated plasma IL-2 concentrations were reported as fold-change over background levels in control, untreated blood samples that were also cultured for 24h. BASTA detected C-peptide responses in >60% (12 of 17) of samples from subjects recently diagnosed with T1D (<100 days), but only in 1 of 8 HLA-matched, non-T1D subjects with no family history of T1D (Mann-Whitney, *P* = 0.0008) (Fig. 1D). This corresponded to an area under the receiver-operator characteristic curve (AUROC) of 0.89 (*P* = 0.0015) for BASTA and 0.52 (*P* = 0.8676) for the CFSE-based Proliferation Assay. Furthermore, the CFSE assay showed lower signal per sample (expressed as the cell division index, CDI [17]) than BASTA (Wilcoxon matched-pairs signed rank test, *P* = 0.04) (Fig. 1F). A higher proportion of samples from non-diabetic subjects had positive responses to C-peptide (CDI>3.0) in the CFSE assay than with BASTA (Fig. 1G). We conclude that BASTA is more sensitive and specific for detecting full-length C-peptide specific responses in T1D than the CFSE-based Proliferation Assay.

**Fig. 1.**
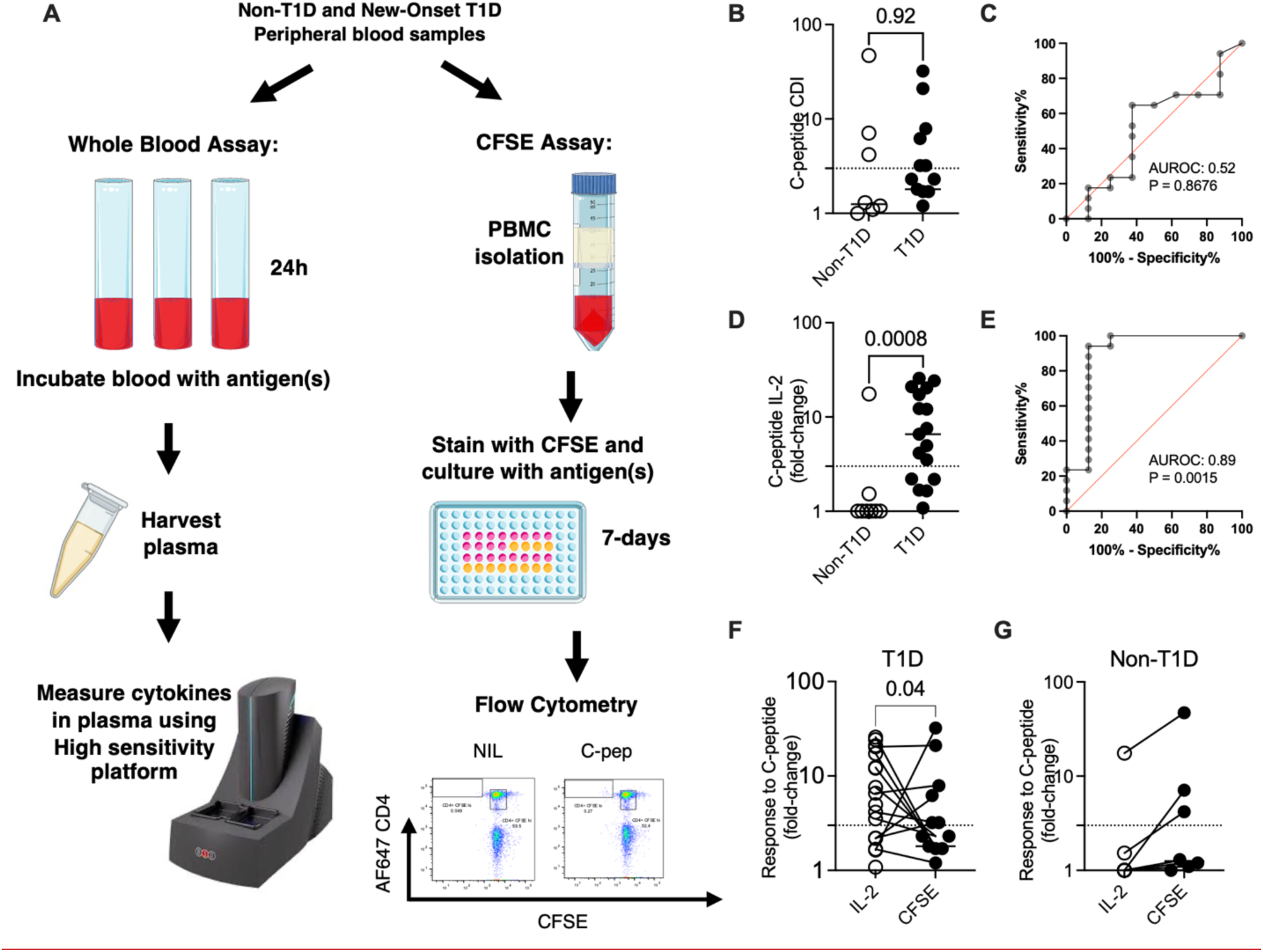
IL-2 secretion in response to C-peptide: a comparison of BASTA and the CFSE-based proliferation assay. (A) Scheme for sampling peripheral blood from subjects with new-onset T1D and non-diabetics for either PBMC isolation for CFSE-based proliferation assay, or direct stimulation with peptide and high sensitivity measurement of IL-2 in plasma. (B) Cell Division Index (CDI) measuring 7-day proliferation of CD4^+^ T cells in response to full-length C-peptide in PBMC samples processed in parallel from 18 donors with newly diagnosed T1D (less than 3 months prior to assay) or 8 HLA-matched (DR4-DQ8, or DR3-DQ2) non-diabetic subjects with 10µM C-peptide, represented as fold-change to background proliferation levels in untreated PBMCs. Dotted line represents 3-fold over background. (C) Receiver operator characteristic (ROC) curve comparing the ability of (B) to distinguish samples from donors with new-onset T1D from those without, determined via the area under the Receiver Operating Characteristic (AUROC). (D) IL-2 levels in plasma after 24h incubation of whole-blood samples from the same participants and blood draw as (A), as a fold-change relative to untreated blood. (E) Receiver Operating Characteristic (ROC) and AUROC calculated from data in (D). (F) Direct paired comparison of C-peptide-induced IL-2 release versus CDI for newly diagnosed diabetic subject, or (G) nondiabetics. Medians are shown for each condition.

### Optimizing BASTA for detecting T-cell responses to preproinsulin

Next, we tested 17 overlapping 18mer peptides which spanned the entire length of human preproinsulin (PPI). These peptides were combined into eight pools of two or three peptides (Fig. 2A). While responses were detected in blood samples from subjects with T1D from across the entire sequence of preproinsulin, the most T1D-specific responses in this cohort were detected to pools 1, 3, and 8, corresponding to the signal peptide, B-chain, and A-chain of preproinsulin (T1D n = 18, non-T1D n = 9) (Fig. 2A).

**Fig. 2.**
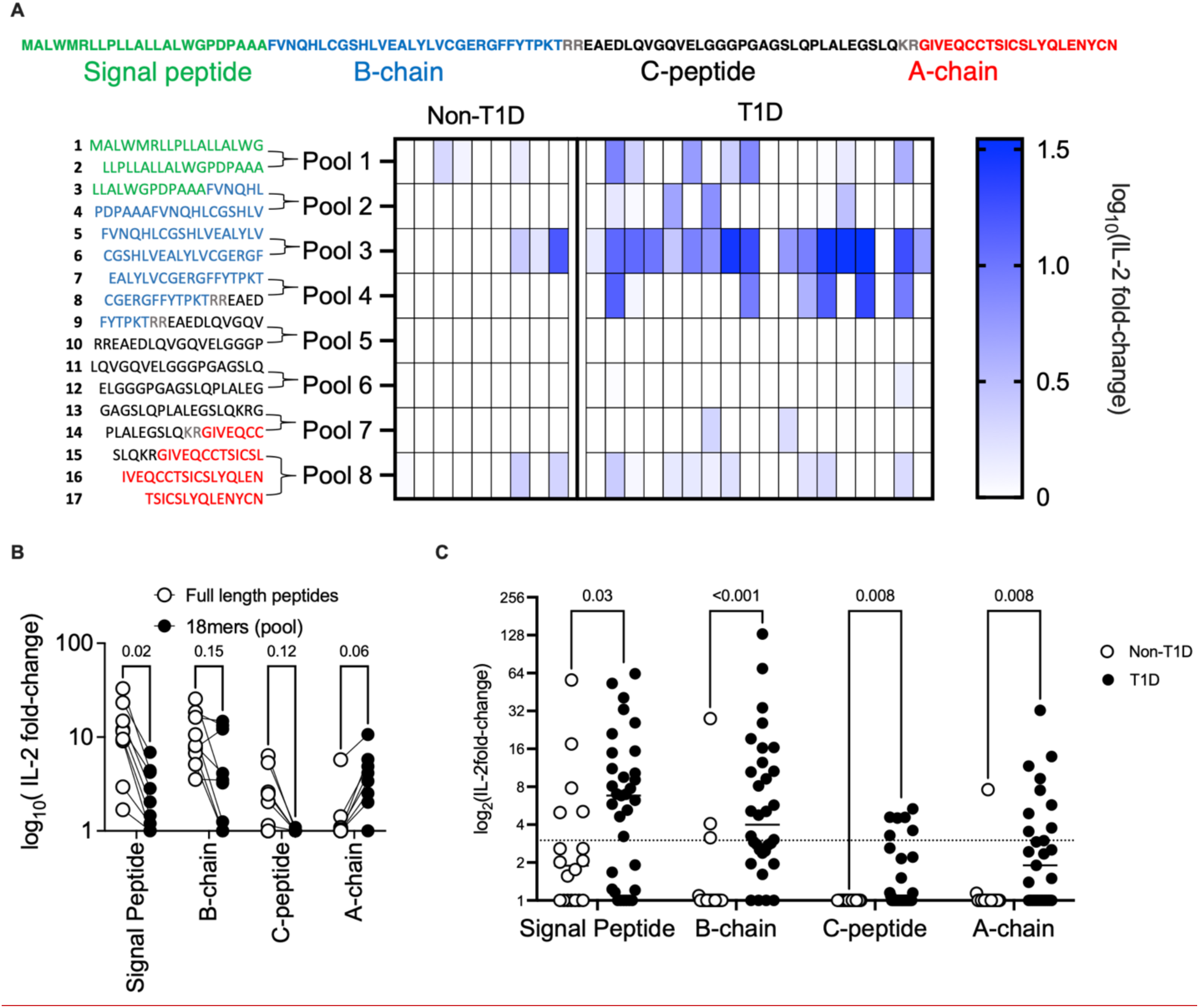
T1D-association of IL-2 responses to peptides across preproinsulin. (A) 17 overlapping 14-18mer synthetic peptides from regions across the length of preproinsulin (PPI) were combined into 8 pools. *Heatmap:* IL-2 responses from whole blood of 9 non-diabetic subjects compared to 18 subjects with established T1D stimulated with 8 peptide pools corresponding to preproinsulin regions across Signal Peptide (green), B-chain (blue), C-peptide (black) or A-chain (red)(junctions in grey not included). (B) Comparison of IL-2 responses to peptide pools in (A) to those induced by individual full-length peptides corresponding to the same preproinsulin region (Signal peptide, B-chain, C-peptide, or A-chain) by stimulated whole blood samples from 6 T1D subjects across a range of time points since diagnosis. (C) Responses to four individual full-length peptides from the four regions of preproinsulin from 19 non-diabetic subjects or 33 diabetic subjects across a range of time points since diagnosis. Dotted line represents 3-fold over background. Horizontal lines depict medians.

Previously, we found that full-length (30 amino acid long) C-peptide (PI_33-63_) was a more potent agonist of several primary CD4^+^ T-cell clones than 18mer peptides incorporating the cognate epitope [13]. To determine the effect of peptide length on T-cell responses measured using BASTA, we compared pools of overlapping18mer peptides to single longer peptide (referred to as ‘full-length’) which covered the sequence of four regions of preproinsulin: signal peptide, B-chain, C-peptide and A-chain (Fig. 2B) (peptide sequences listed in Table S3). Full-length signal peptide stimulated stronger responses in samples from people with T1D (n=9) than overlapping 18mers, each at the same molar concentration (Wilcoxon multi-test, adjusted *P* = 0.02). Consistent with our previous findings, 56% (5 of 9) samples responded to full-length C-peptide, but none responded to the corresponding pooled 18mers (0 of 9). In contrast, responses to insulin B-chain were unaffected by peptide length and responses to full-length A-chain were diminished compared to 18mers (Wilcoxon multi-test, adjusted *P* = 0.06). A concentration of 10.0µM C-peptide induced highly disease-specific responses. Titration experiments revealed that 2.0µM preproinsulin signal peptide and insulin B-chain increased the T1D specificity of responses (Fig. S2A), whereas 10.0μM was optimal insulin A-chain.

The presence of cysteines in the peptides can impact their immunogenicity [27] and stability. Full-length insulin A-chain is very difficult to synthesize and work with due to its insolubility and reactivity [34], which may have contributed to its relatively poor performance in these assays. To find more stable variant peptides we evaluated the effect of substituting cysteine residues with serine in insulin B-chain and A-chain. Serine substitution for cysteine had no apparent effect on responses to B-chain for samples, but as expected (see [27, 28]), destroyed recognition of the A-chain peptide (Fig S2B). To address the issues with handling the full-length insulin A-chain peptide we introduced an isoacyl bond between Thr8 and Ser9 [35]. The isoacyl bond both increases the solubility of the peptide and suppresses aggregation, but spontaneously isomerizes to native A-chain at physiological pH [36, 37]. Full-length insulin A-chain elicited responses from all three test T1D samples (Fig. S2B).

In a pediatric clinical setting, the blood volume required for a test is often limiting. To determine the minimum volume of blood required for the assay responses to preproinsulin peptides from 1.0, 0.5 or 0.25mL of blood were compared, maintaining the same peptide concentration. There was no decrease in IL-2 response to all four preproinsulin peptides using 0.5mL compared to 1.0mL (Fig. S2C-D) of blood from three individuals. However, reducing the volume to 0.25mL significantly increased the inter-assay variability (Wilcoxon matched pairs signed rank test, *P* = 0.0494) (Fig. S2E). Hence, we chose 0.5mL blood per antigen treatment for future assays. We also compared preproinsulin induced IL-2 responses after culture in two different vessels: 1.0mL polypropylene cryovials (Falcon), or 5.0 mL round-bottom polypropylene capped tubes (SPL Life Sciences) (Fig. S2G). Responses were not significantly different between the different tubes and both gave similar percentage coefficient of variation (%CV). We chose to use 5.0mL capped tubes for handling convenience. In summary, the optimized BASTA format is: full-length peptides at a final concentration of 2.0μM for the preproinsulin signal peptide and insulin B-chain, or 10.0μM for C-peptide and insulin A-chain (as isoacyl A-chain), delivered as concentrated peptide stock in minimal solvent to 0.5mL whole blood per antigen treatment in a capped 5.0mL polypropylene tube.

### T1D-associated IL-2 responses to proinsulin peptides

Individual peptides with sequences corresponding to all four regions of preproinsulin (signal sequence, B-chain, C-peptide and A-chain) stimulated increased IL-2 responses in 33 subjects with T1D compared to 19 non-T1D samples (Multiple Mann-Whitney tests, Signal peptide: adj. *P* = 0.03, B-chain: adj. *P* = 0.00008, C-peptide adj. *P* = 0.008, A-chain adj. *P* = 0.008) (Fig. 2C). With the aim of further minimizing volume of blood required, we tested responses against a pool of all four full-length preproinsulin peptides (Signal peptide, B-chain, C-peptide, isoacyl-A-chain). Responses from 11 adult participants without T1D, including 36% (4/11) with T1D-associated high-risk HLA-DR3-DQ2 and HLA-DR4-DQ8 haplotypes, were compared to 20 individuals with T1D (disease duration ranging from the day of diagnosis to 12 years). IL-2 responses to the preproinsulin peptides were clearly elevated in samples from individuals with T1D (Mann-Whitney, *P* <0.0001). Responses to the preproinsulin pool were very effective at distinguishing subjects with T1D from those without T1D, giving an AUROC of 0.93 (*P* < 0.0001) (Fig. 3A). A majority of T1D subjects in this cohort had longstanding T1D, which contributed to the low proportion of C-peptide responders (Fig. S3A) (as previously observed in [13]).

**Fig. 3.**
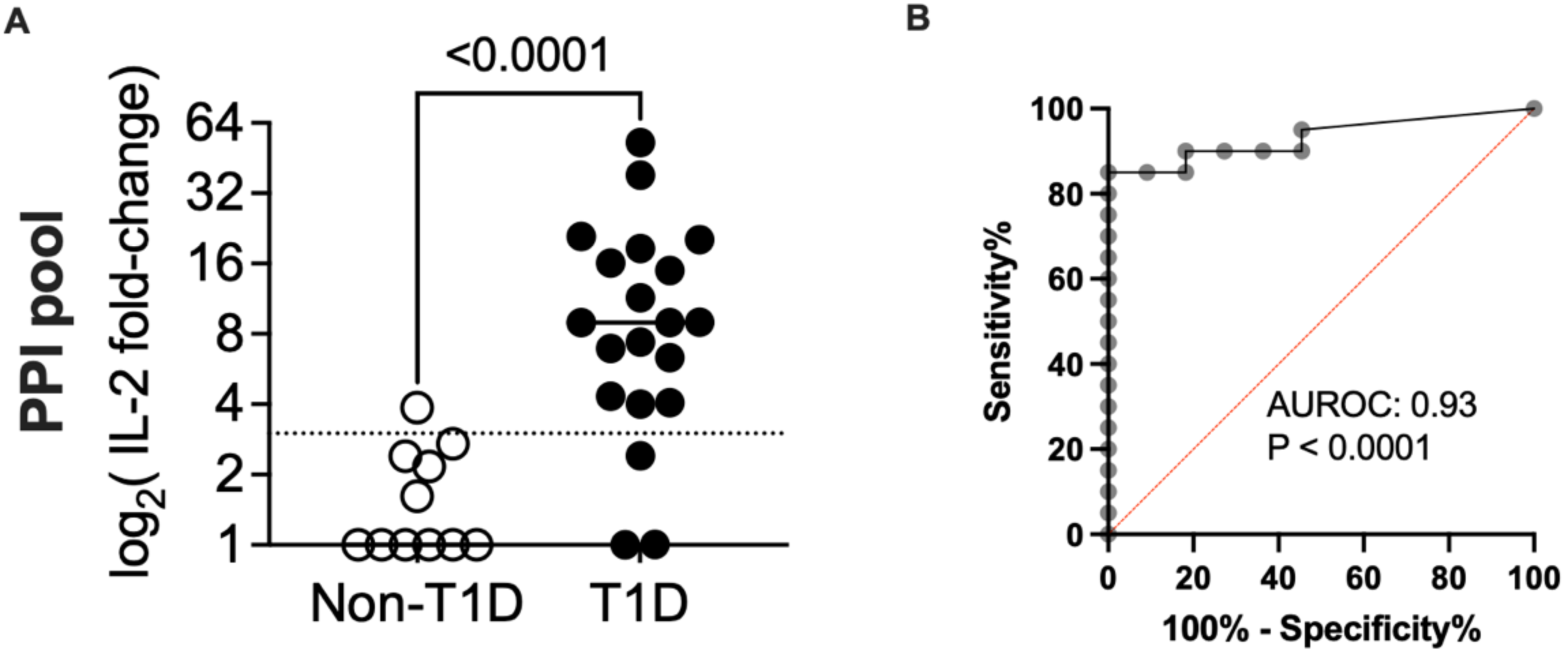
Optimized assay format reveals stability of disease-specific CD4^+^ T-cell responses to preproinsulin. (A) Responses to a pool of peptides from all four regions of preproinsulin as measured by IL-2 released in whole blood samples from non-diabetic subjects or diabetic subjects across a range of time points since diagnosis. (B) Receiver Operating Characteristic (ROC) curve for prediction of T1D status based on levels of IL-2 as measured in (A), area under the curve (AUC): 0.93. Dotted line represents 3-fold over background. Horizontal lines depict medians.

We next evaluated the inter-and intra-assay variability of BASTA using individual preproinsulin peptides and a pool of all four preproinsulin peptides. The intra-assay variability was low (Fig. 4A). In samples from four subjects with established T1D, duplicate stimulations of blood from the same draw using the preproinsulin pool in separate stimulation tubes had a mean CV of 13.7%, and duplicate wells of plasma in the IL-2 assay had a mean CV of 3.7%. While this provides evidence of strong reproducibility, it indicates that individual blood aliquots and their stimulation are a greater source of experimental variation than the IL-2 measurements (Wilcoxon signed-rank test, *P* = 0.0076) (Fig. 4A). The inter-assay variability was determined by testing samples from four individuals with longstanding T1D (T1D#1-4) on four occasions 1-3 weeks apart. Triplicate whole-blood cultures were set up for each individual preproinsulin antigen, preproinsulin pool, or tetanus toxoid. After collecting the triplicate plasma samples, a single IL-2 measurement was made from each. All individuals consistently exhibited responses to the preproinsulin pool (Fig. 4B). Inter-assay variability for responses to the preproinsulin pool had a mean CV of 31.7-54.7% (Fig. S4B). For individual antigens, the inter-assay variability was lowest for responses to tetanus toxoid (mean CV: 11.72%) and highest for responses to preproinsulin signal peptide (mean CV: 53.9%) (Fig. S4C).

**Fig. 4.**
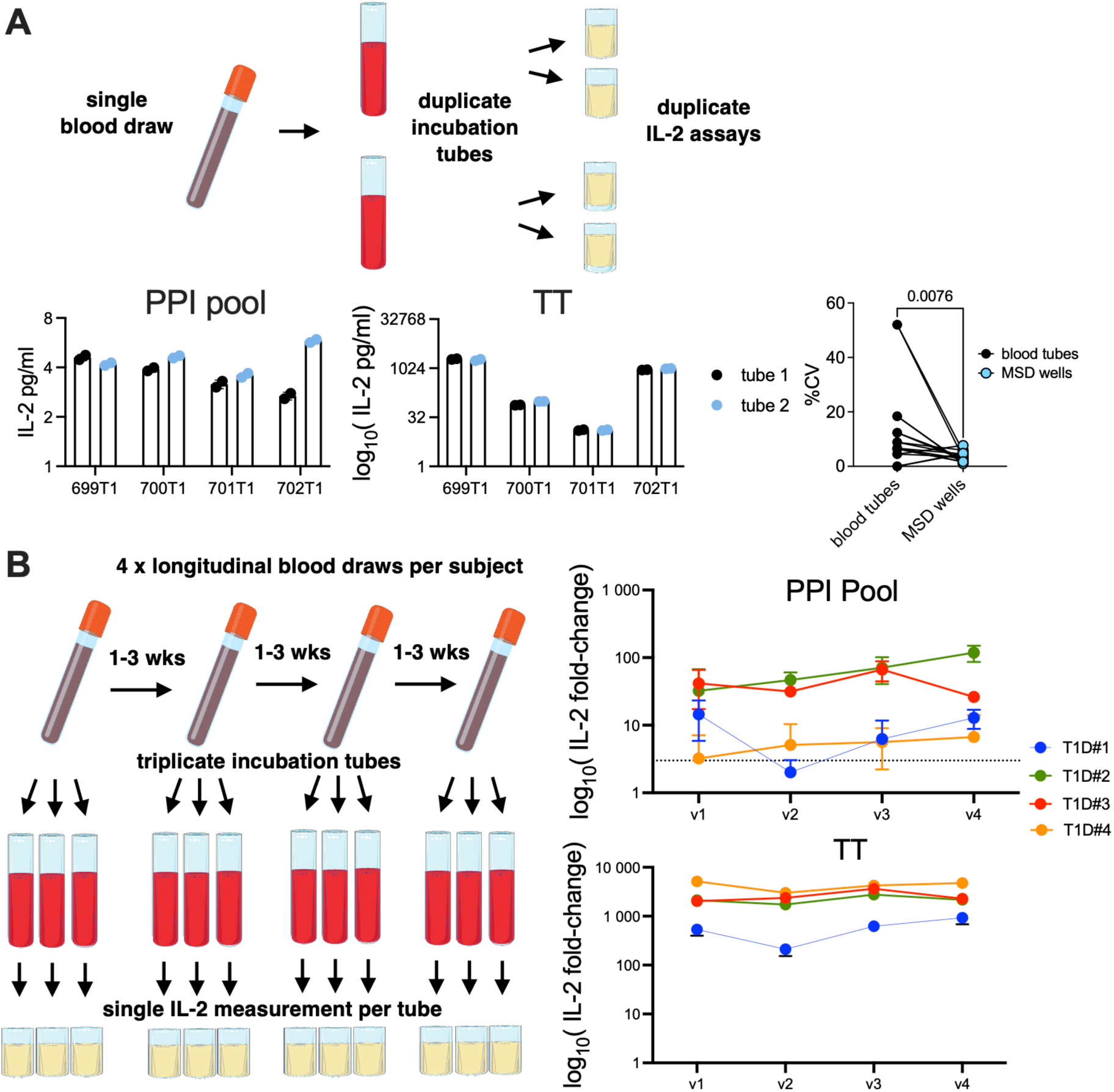
Robustness of preproinsulin-induced IL-2 responses from whole blood across stimulation replicates and stability across multiple longitudinal measurements. **(A)** Assessment of intra-assay variability between duplicate stimulation tubes (black versus blue) from the same blood draw, and duplicate IL-2 measurements of plasma (individual dots). *Right:* Comparison of %CV as variance between stimulation replicates (blood tubes) versus IL-2 measurements from the same plasma (MSD plate wells). **(B)** Repeat measures of IL-2 responses to individual or pooled peptides from all regions of preproinsulin (Signal peptide, B-chain, C-peptide, A-chain, or all four together), or tetanus toxoid, from 4 subjects with established T1D (T1D#1-4), measured in triplicate on four occasions with intervals of 1-3 weeks between blood draws.

### Whole Blood IL-2 release assay measures antigen-specific CD4^+^ memory T-cell responses

We then sought to determine the cellular source of the IL-2 detected in the whole blood assay in responses to antigenic stimulation. First, we investigated the requirement for HLA class II by adding HLA-class II blocking mAbs to the whole blood culture. IL-2 release was modestly inhibited in the presence of blocking mAb specific for HLA-DR, -DP and -DQ, but almost completed inhibited in the presence of all three mAbs (One-way ANOVA with Holm-Sidak multiple-comparisons test, adj. *P* = 0.0003) (Fig. 5A). Similar trends were observed from tetanus toxoid stimulations (*P* = 0.0775) (Fig. S5B). In contrast, an anti-HLA class-I mAb (W6/32), of the same isotype (murine IgG2a) did not attenuate the IL-2 responses (Fig. S5A). Next, we directly identified the IL-2 producing cells using IL-2 surface capture and flow cytometry. In response to a pool of preproinsulin peptides, or tetanus toxoid, a distinct population of IL-2^+^ CD4^+^ T-cells was seen (Fig. 5B). The number of CD4^+^, IL-2^+^ T-cells recovered using the surface IL-2 capture assay was strongly correlated with IL-2 concentrations from a whole blood assay run in parallel (Pearson *r* = 0.93, *P* < 0.0001) (Fig. 5C). Phenotypic analysis of the IL-2 producing cells (Fig. S5C) revealed that after preproinsulin peptide stimulation, the CD4^+^, IL-2^+^ T cell subset contained significantly fewer naïve cells (CD45RO^-^, CCR7^+^) and more CD45RO^+^ memory cells than the CD4^+^, IL-2^-^ subset (multiple paired T-test, adj. *P* = 0.03 for T_naive_, adj. *P* = 0.02 for T_mem_) (Fig. 5D). The CD4^+^, IL-2^+^ T cell subset shows a trend towards being enriched in CCR7^-^ effector/effector memory T cells (adj. *P* = 0.09). Furthermore, CD69 was found exclusively expressed on CD4^+^, IL-2^+^ T cells (Mann-Whitney, *P* = 0.008) (Fig. 5E). Hence, we conclude that BASTA measures memory/effector CD4^+^ T-cell responses to β-cell antigens and tetanus toxoid.

**Fig. 5.**
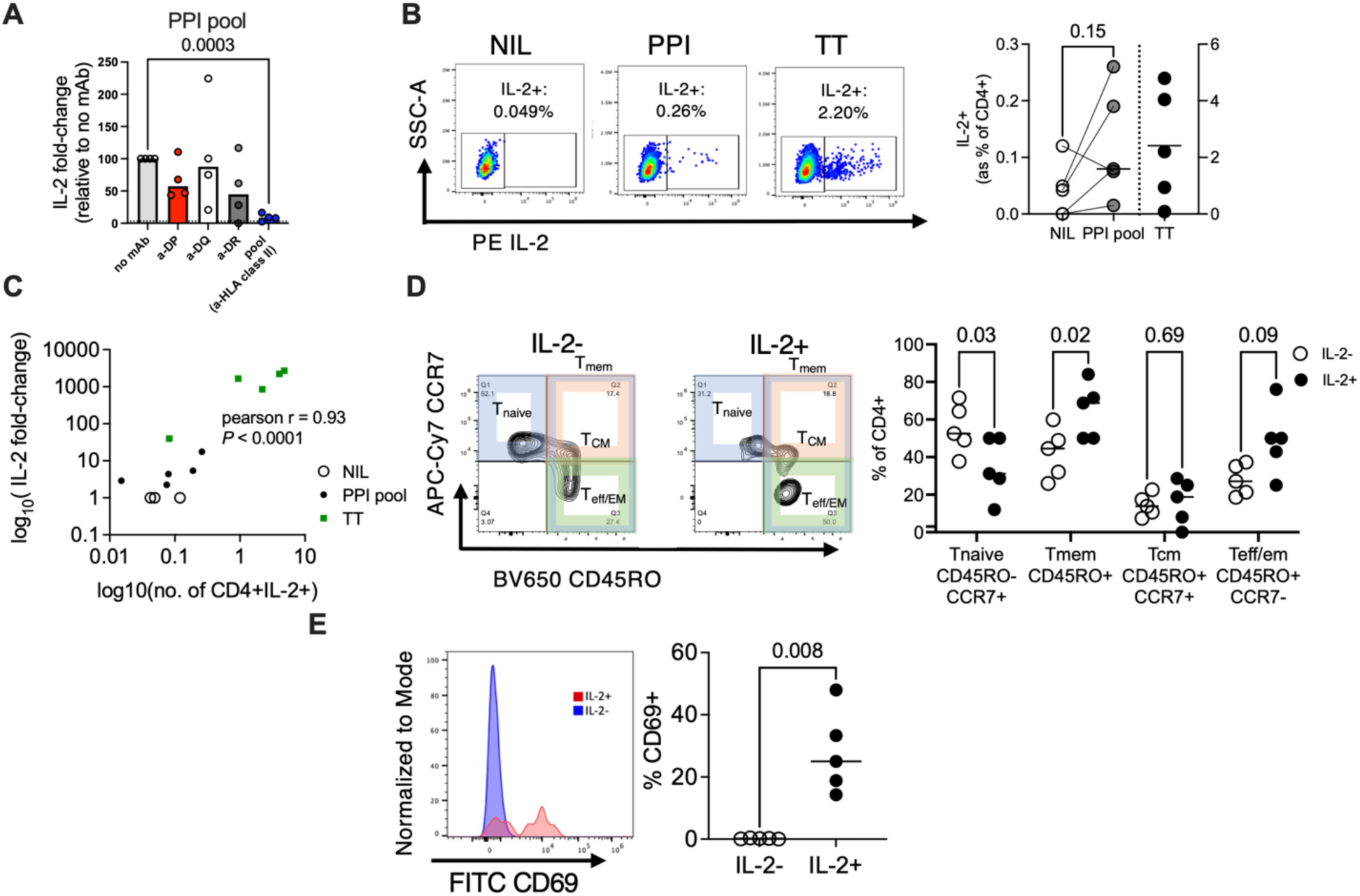
Antigen-reactivated CD4^+^ T cells are the major IL-2 secreting population in whole blood. **(A)** The effect of HLA-DP, -DQ, -DR, or a combination of all three, blocking mAbs on PPI-induced IL-2 release relative to PPI-stimulated blood alone (bars show median) (n = 4 x T1D). **(B)** FACS plots and proportion of IL-2^+^ cells as detected by IL-2 capture assay in CD4^+^ T cells isolated from untreated, PPI-or tetanus toxoid-stimulated whole blood following magnetic enrichment of IL-2 secretors. **(C)** Correlation between secreted IL-2 as detected via MSD immunoassay in plasma compared to the percentage CD4^+^, IL-2^+^ detected by the IL-2 capture assay. **(D)** Proportions of T_naive_ (CD45RO^-^, CCR7^+^), T_mem_ (total CD45RO+), T_CM_ (CD45RO^+^, CCR7^+^) and T_EFF/EM_ (CD45RO^+^, CCR7^-^) within the IL-2^+^ versus IL-2^-^, CD4^+^ T-cells in a PPI-stimulated whole blood assay as detected by capture assay. **(E)** Representative histograms and proportion of CD69^+^ within IL-2^-^ and IL-2^+^ CD4^+^ populations as in D. Lines at median are shown. B-D show data from 5 individual subjects with T1D.

### BASTA measures T1D-specific CD4^+^ T-cell responses in children

Up to this point we had largely used HLA-matched adult donors for the non-T1D control samples. To determine if the age of the subject impacted CD4^+^ T-cell responses to preproinsulin, we validated the assay by measuring responses to preproinsulin peptides in a cross-sectional cohort of children and adolescents, ages 3-to 20-years-old (as per scheme Fig. 6A, see also Table S2 for participant details). Test subjects included participants without T1D (“low-risk”), recruited from the community (n = 10) as well as children at-risk for developing T1D because they have a first-degree relative (parent or sibling) with T1D. Samples for the whole blood assays were taken at a single time point in subjects greater than 3 years of age. These at-risk participants were further stratified into two groups: (i) negative for islet autoantibodies (“AAB-”) (n = 14), or (ii) positive for one or more islet autoantibody (“AAB+”) (n = 8). Subjects with a clinical T1D diagnosis were also recruited and divided into two groups: (i) pediatric participants within 100 days of diagnosis with T1D (“New-Onset T1D”) (n = 12), or (ii) those with established T1D, > 100 days post diagnosis (“Established T1D”) (n = 20). In contrast to the test cohort using adult non-diabetic samples (in Fig. 3A), whole blood from young non-T1D pediatric subjects in the AAB^-^ and AAB^+^ groups gave IL-2 responses to the pool of four preproinsulin peptides similar to those from T1D participants (Fig. 6B). As a result, the preproinsulin pool did not distinguish samples based on T1D status (AUROC = 0.55, *P* = 0.5182) (Fig. 6C), with no increase in AUROC by comparing non-T1D samples with only New-Onset T1D (Fig. 6D). In marked contrast, responses to full-length C-peptide were clearly associated with T1D (Fig. 6D). These largely distinguished non-T1D from T1D samples, although lower prevalence of C-peptide responders in established T1D participants lowered the overall AUROC (AUROC = 0.77, *P* = 0.0002) (Fig. 6E). CD4^+^ T-cell responses to C-peptide detected by BASTA were markedly different between new-onset T1D and subjects without T1D (AUROC = 0.86, *P* = 0.0003) (Fig. 6F). When CD4^+^ T-cell responses to individual preproinsulin peptides were evaluated, many pediatric non-diabetic subjects had detectable responses to the preproinsulin signal peptide and insulin B-chain. There was no difference between autoantibody positive (AAB^+^) and negative (AAB^-^) non-diabetic children with a family history of T1D (Fig. S6). In contrast, non-T1D subjects, with no family history of T1D, showed significantly lower responses to insulin B-chain compared to the other groups, notably the AAB^-^ non-T1D children with a first-degree relative with T1D (Kruskal-Wallis using Dunn’s post-hoc test, adj. *P*=0.04) (Fig. S6B). A similar trend is reflected in the differential responses of these groups to the preproinsulin pool (*P* = 0.06) (Fig. S6A). As in the initial test cohort (Fig. 1D), responses to C-peptide above 3-fold over background are highly specific for individuals with T1D and most prevalent in New-Onset T1D (Kruskal-Wallis using Dunn’s post-hoc test comparing non-T1D to New-Onset T1D, adj. *P* = 0.01) (Fig. S6D). Responses to the isoacyl insulin A-chain were low in all groups (Fig. S6E). IL-10 secretion was minimal in response to all stimuli for all subject groups (Fig. S7A). When stratifying for HLA-type, individuals with T1D who are heterozygous for high-risk HLA-DR3-DQ2 and HLA-DR4-DQ8 haplotype showed the strongest IL-2 responses to full-length C-peptide (Kruskal-Wallis using Dunn’s post-hoc test comparing non-T1D X/X to T1D DQ2/DQ8 [adj. *P* = 0.0026]) (Fig. S7B). Upon further stratification, responses to C-peptide were also indeed higher in samples from at-risk, due to their family history, pediatric donors if positive for multiple islet autoantibodies, compared to those positive for a single autoantibody (Kruskal-Wallis using Dunn’s post-hoc test, adj. *P* = 0.02) (Fig. S7C). This was specific for C-peptide, as increased IL-2 from multi-antibody-positive individuals was not observed in response to any other preproinsulin peptide (Fig. S7C). We conclude that full-length C-peptide, but not the other preproinsulin peptides tested, stimulates T1D associated CD4^+^ T-cell responses in young people which we can be measured with BASTA.

**Fig. 6.**
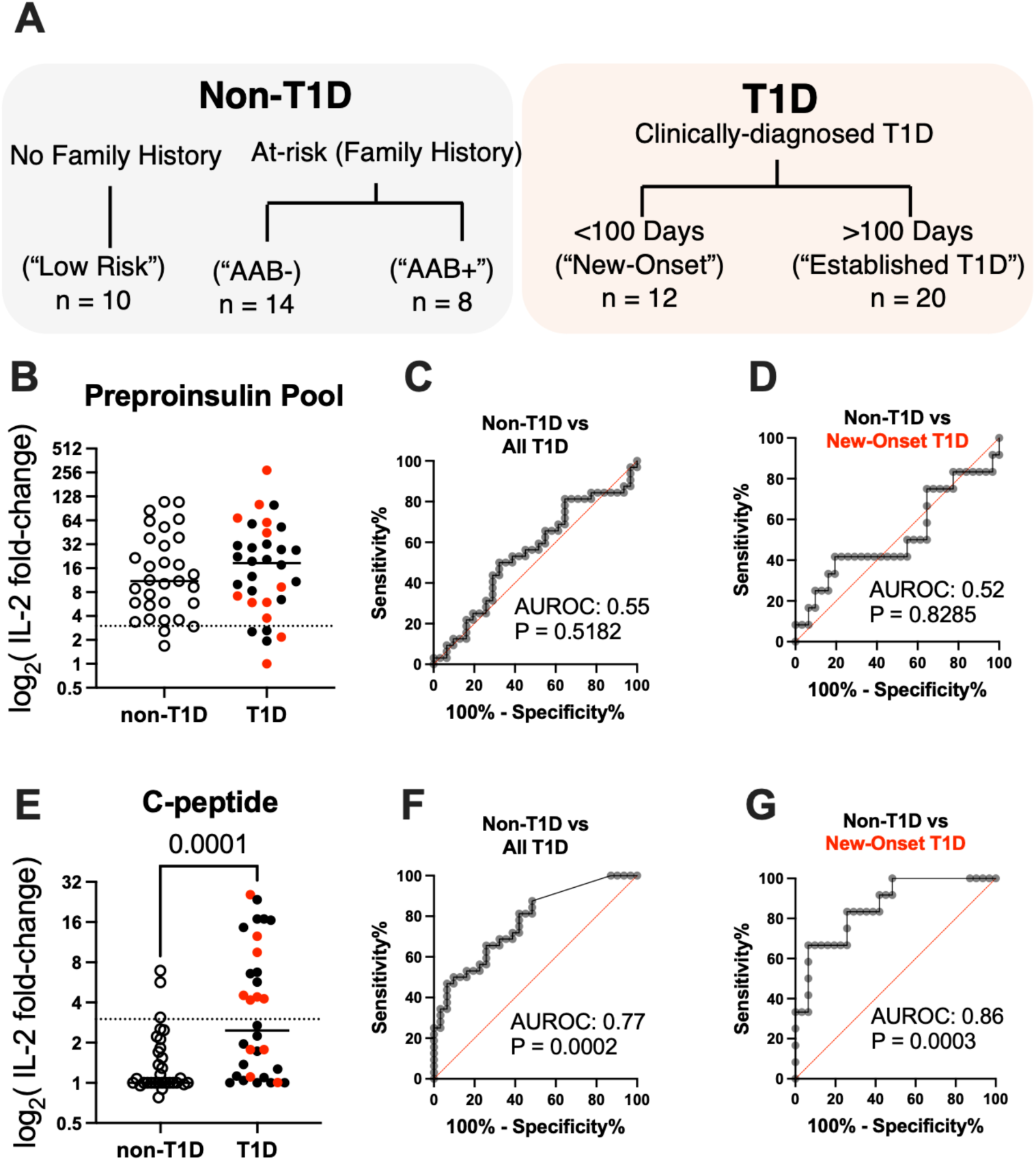
Cross-sectional analysis of IL-2 responses to combined and individual PPI-peptides in non-diabetics, or subjects with T1D using the optimized Whole Blood Assay. **(A)** Individuals were categorized as either non-T1D (participants without T1D and no family history (“low risk”), or AAB-and AAB^+^ pediatric subjects with a first-degree relative with T1D), or T1D, being clinically diagnosed <100 days (“New-Onset”) or >100 days (“Established T1D”) prior to sampling. **(B)** Comparison of IL-2 release in response to the preproinsulin peptide pool from non-T1D and T1D samples (red circle = New-Onset T1D). **(C)** ROC and AUROC calculated from data in (B), to distinguish non-T1D from all T1D samples, with **(D)** comparing non-T1D to only New-Onset T1D. **(E)** IL-2 responses to C-peptide, with corresponding ROC and AUROC in **(F)** using C-peptide responses non-T1D from all T1D or **(G)** comparing only New-Onset T1D to non-T1D. Dotted line represents 3-fold over background. Lines at median are shown.

## DISCUSSION

Here we describe BASTA, a simple whole blood assay that is suitable for the routine clinical measurement of β-cell antigen-specific CD4^+^ memory T-cell responses. This assay overcomes many of the obstacles inherent in assays for CD4^+^ T-cell autoimmunity against β-cell antigens. The major advantages of this assay include: (i) it requires minimal blood, as low as 1.0 ml (for two 0.5ml replicates) per antigen treatment, (ii) it is very simple to perform, (iii) it has a short 24h culture period, (iv) plasma samples can be stored, shipped and analyzed in batches, (v) the platform can be easily adapted to different antigens and peptide epitope combinations. We anticipate that this assay will be very useful for the monitoring of changes in β-cell antigen-specific memory CD4^+^ T-cell function in clinical trials and other clinical research settings.

Responses to full-length C-peptide (PI_33-63_) consistently show highly specific and sensitive association with new-onset T1D. This is consistent with our earlier reports showing that full-length C-peptide is a central antigen in human T1D [12, 13]. Much other previous work has focused on T-cell responses to insulin B_9-23_ in association with T1D [38–40], however peripheral blood responses to the B-chain region of preproinsulin are often detected in non-diabetic subjects [40]. BASTA responses to the insulin B-chain were frequently detected in individuals without T1D, including from many non-diabetic pediatric donors with a parent with T1D. Interestingly, these were largely absent from the non-T1D pediatric subjects with no family history of T1D. Furthermore, the preproinsulin signal peptide was a strong agonist of IL-2 responses from both people with and without T1D, adult and pediatric subjects alike. In contrast, only the C-peptide and, to a lesser degree, insulin A-chain showed specificity for T1D (Fig. 2C, Fig. 6). Although responses to insulin A-chain have been well characterized [27], response to full-length A-chain peptide were relatively weak and detected rarely. Pooling of preproinsulin peptides did show advantages in greatly enhancing the proportion of responders (as in Fig. S4A) and the absolute IL-2 concentrations, but at the cost of reduced T1D specificity in younger donors (Fig. 6B, C). This suggests that pooling peptides will be a useful strategy for increasing the breadth of T-cell responses measured, but that the composition of this peptide pool needs to be carefully evaluated.

In response to preproinsulin peptides, CD4^+^ T-cell effector cells are the predominant source of IL-2. We found little evidence for IL-2 production by CD8^+^ T-cells, or other cell types. This is not surprising since we have used relatively long peptides which we expect to favor HLA class II restricted, CD4^+^ T-cell responses and a relatively short culture period, which favors activation of memory T cells. An increase in IL-10 secretion, in response to preproinsulin and other islet autoantigens has been reported in non-diabetic siblings of children with T1D [41]. IL-10 was interpreted to be a surrogate for regulatory T cells. However, in our hands, we did not detect any evidence that IL-10 is useful for measuring regulatory T-cell responses. Although IFNγ is commonly measured in immunoassays, the relatively high background of IFNγ in cultured whole blood makes this cytokine unsuitable for this application. IL-2 signals were reproducible across assays, and showed comparable variability (%CV) to previously described T-cell assays for β-cell antigens using tetramers (mean CV 14-23%) [42, 43] or the CFSE-based Proliferation Assay (mean CV 30-53%.) [44], presumably due to the low precursor frequency of β-cell antigen specific CD4^+^ T cell in peripheral blood.

The preliminary assay validation and optimization used samples from adults without T1D (>20 years of age) as negative controls, compared to T1D samples recruited from a pediatric (<18 years of age) endocrinology clinic. However, preproinsulin signal peptide and insulin B-chain later showed less T1D specificity when used to analyze samples from T1D children compared to age-matched negative controls. Most strikingly, while responses to the preproinsulin pool from non-T1D adults were low or absent (Fig. 3A), responses to this pool from non-T1D children were similar to those of age-matched children with T1D (Fig. 6B). This could indicate dynamics of beta cell antigen-specific T-cell reactivity that change with age, or more simply general changes to reactivity of T-cells. Our test cohort included children with and without islet autoantibodies. While the overall presence or absence of islet autoantibodies did not give any clear association between CD4^+^ T-cell responses to preproinsulin peptides (Fig. S6A-E), further subset analysis revealed that multi-antibody positive children had stronger C-peptide responses compared to single antibody positive children, in accordance with the heightened T1D risk in this subgroup [4]. This further indicates a potential role for full-length C-peptide-directed CD4^+^ T-cell responses as a biomarker of stage 1 or 2 T1D.

There are some limitations of our work. We have focused on a set of preproinsulin-derived peptides and pools of preproinsulin peptides. While it is clear from our data that C-peptide stimulates T1D specific CD4^+^ T-cell responses, we are actively evaluating other antigens with the aim of curating a pool of validated peptides which stimulate T1D specific responses. Each new candidate peptide needs to be carefully evaluated to identify those that stimulate T1D specific CD4^+^ T-cell responses. As our data shows, many ‘self’ peptides, such as preproinsulin signal peptide, stimulate CD4^+^ T-cell response in individuals without β-cell autoimmunity. One of the advantages of this assay format is that suitable β-cell protein derived peptides can easily be added to the assay as they are identified. The BASTA assay will be improved by further optimization. Specifically, we will focus on reducing the inter-assay variability. We have not undertaken an extensive evaluation of the stability and solubility of different peptides used in the assay. Based on previous results [33], all blood stimulations began within 3 hours of blood draw, but this was not re-examined for responses to islet peptides. These parameters will need to be optimized to increase the robustness and durability of the assay. IL-2 levels in control, unstimulated whole blood samples were often well below the calculated lower limit of detection (LLOD) for the MSD V-plex technology used, and as such, using a format with higher sensitivity (such as the MSD S-plex, which can measure down to fg/ml) could further enhance the sensitivity of this assay. These steps for further development will be important for using BASTA to investigate changes in β-cell antigen specific autoimmunity. Currently, we have not attempted to use BASTA to predict when an individual will progress to Stage 3 T1D. However, eventually BASTA may allow more accurate prediction of the time an individual will progress to Stage 3 T1D and require insulin therapy. At this stage, we have not used BASTA to monitor changes in β-cell antigen specific CD4+ T cell responses in the context of a clinical trial. Nonetheless, our data show that BASTA is both sufficiently sensitive and specific for measuring T1D specific CD4^+^ T-cell responses and we anticipate that it will be find may applications in the clinical and research settings.

In conclusion, T1D specific CD4^+^ T-cell responses to full-length C-peptide can be readily measured using a simple whole blood-based assay. Due to its simplicity and robustness, we believe BASTA will be a useful and powerful tool for both dissecting autoimmune CD4^+^ T-cell responses and monitoring changes in clinical trials and community settings. Our work paves the way for the development of a long-sought after T-cell assay that is suitable for routine clinical monitoring of CD4^+^ T-cell responses in people with, or at risk of T1D.

## MATERIALS AND METHODS

### Study Design

This study was conducted in two phases. For the initial phase, patients with T1D were recruited during regular consultations at The Royal Children’s Hospital (Victoria, Australia) endocrinology clinic or at baseline sampling prior to entry into the BANDIT trial [45]. Non-diabetic control samples were recruited through the SVI Living Biobank. For the validation phase, additional samples were obtained from research participants attending Royal Melbourne Hospital. Ethical approval was given by St Vincent’s Hospital HREC (Approval Numbers: HREC-A 161.15 and HREC-A 135/08) and Southern Health (Royal Children’s Hospital, approval number: 12185B). Further approval was given by Melbourne Health (Approval number: 2009.026). All participants provided written informed consent. Participants are without T1D or diagnosed according to American Diabetes Association criteria. HLA typing of donor samples was provided by Victorian Transplantation and Immunogenetics Services (Victoria, Australia). Blood donors were classed based on the presence of HLA-DR3-DQ2 (HLA-DQB1*02:01, HLA-DQA1*05:01) or HLA-DR4-DQ8 (HLA-DQB1*03:02, HLA-DQA1*03:01) haplotypes with high risk for T1D. Donors possessing the HLA-DQA1*03:03 variant were also included within the HLA-DQ8 haplotype (as established in [46]).

### Peptides and Stimuli

Preproinsulin peptides at > 90% purity (supplied by Genscript except for isoacyl A-chain, which was prepared in-house) were dissolved in DMSO, or acetonitrile/water with or without the addition of 0.5% acetic acid, or 100mM ammonium bicarbonate, to a concentration of 5.0mM, aliquoted and stored at ^-^80°C. Pre-dissolution of isoacyl A-chain in acidic conditions prevented base-mediated isomerization. A full list of peptides is shown in Table S3. Initial experiments used commercially available CEFX (JPT Peptide Technologies, Germany), a pool of peptide epitopes from pathogens known to stimulate CD4^+^ T-cell responses, at 1µg/ml. Tetanus toxoid was used at 0.33Lfu/ml and supplied by Statens Serum Institut, Copenhagen, Denmark.

### The β-cell antigen specific T-cell assay -BASTA

Briefly, heparinized peripheral blood was aliquoted within 3 hours of blood draw into sterile, capped, polypropylene tubes containing pre-aliquoted peptides or control stimuli, or similarly prepared cryovials with external-threaded caps. Blood without any additional stimulation, or the peptide solvent alone was an additional negative control. After 24-hour incubation at 37°C with 5% CO2, samples were centrifuged, plasma was harvested, aliquoted and frozen. Cytokines in the plasma were measured in duplicate using an electrochemiluminescence (ECL)-based immunoassay platform (Meso Scale Discovery (MSD), Rockville, MD, USA), using lot-validated 96-well plate format MSD V-plex kits measured on a MESO QuickPlex SQ 120 reader. Data was analysed using the Discovery Workbench software. If below LLOD results were set to the LLOD for each assay run based on the standard curve. Response cut-off was selected as 3-fold over the unstimulated or solvent only control blood cultures.

### The CFSE-Based Proliferation Assay

The CFSE (5,6-carboxylfluorescein diacetate succinimidyl ester) proliferation assays were performed as described previously [17]. Briefly, PBMC from T1D subjects or non T1D HLA matched donors labelled with 0.1mM CFSE (Life Technologies, Carlsbad, CA) were cultured either with: no additional stimulus, or C-peptide (10μM), or tetanus toxoid (0.33LfU/ml). After 7 days of culture the cells were washed in PBS and stained on ice with anti-human CD4-AlexaFluor-647 (clone OKT4, prepared in house). CD4^+^ T-cell proliferation was measured by determining the number of CD4^+^, CFSE^dim^ cells for every 5,000 CD4^+^ CFSE^bright^ cells. The results are presented as a cell division index (CDI) which is the ratio of the number of CD4^+^ cells that have proliferated in the presence of antigen: without antigen [47].

### HLA-blocking

As previously described [28, 47], TCR recognition of pMHC complexes on antigen presenting cells was blocked through the addition of monoclonal antibodies specific for either HLA-DP (clone: B7/21), HLA-DQ (clone: SPV-L3), HLA-DR (clone: L243), the addition of all 3 pooled antibodies, or pan anti-HLA-class I (clone: W6/32, 2µg/ml final concentration). Each was added in combination to either blood with no additional stimulus or peptide/protein stimuli as indicated.

### IL-2 capture

IL-2^+^ cells were enriched for further analysis using a magnetic bead-based IL-2 capture system (Miltenyi) according to manufacturer’s instructions. Briefly, whole blood was stimulated overnight as above, in 5ml volumes using 50ml tubes. After 16h incubation, erythrocytes were lysed (using Red Cell Lysis Buffer, Miltenyi), and cytokine secreting cells identified through further labelling with capture reagents and incubation. Cells were then stained with PE-labelled anti-IL-2 antibodies, and the following panel of antibodies against surface markers: CD3-PerCP (clone: UCHT1, Biolegend), CCR7-APC-Cy7 (clone: G043H7, Biolegend), CD45RO-BV650 (clone: UCHL1, BD biosciences), CD4-AF647 (clone: OKT4, prepared in-house), CD8-V500 (clone: RPA-T8, BD biosciences), CD69-FITC (clone: FN50, BD Biosciences), prior to magnetic enrichment using MS columns.

### Flow cytometry

Flow-cytometry was performed as previously described [13, 17], cells were acquired using the flow cytometer LSRFortessa (BD Biosciences) or Cytek Aurora (Cytek Biosciences) and analyzed in FlowJo 10 (v10 TreeStar). Live cells were gated on by excluding cells positive for propidium iodide (PI).

### Statistical analysis

Statistical tests were performed using GraphPad Prism 10 for MacOS (GraphPad Software, San Diego California USA). Datasets with a normal distribution were analyzed using paired or unpaired two-tailed Student’s t-test. Non-parametric data was assessed using an unpaired two-tailed Mann-Whitney U-test or Wilcoxon matched-pairs signed rank test, corrected for multiple comparisons where appropriate. P-values ≤ 0.05 were considered statistically significant. Analysis between multiple groups was performed using ANOVA or Kruskal-Wallis test for non-parametric data, with Dunn’s post-hoc test for comparison between individual groups. Area under the receiver operator characteristic curve (AUROC) calculated using inbuilt function in Prism. The Coefficient of variation (CV) was calculated as the ratio of the standard deviation to the mean, expressed as a percentage (that is, %CV = SD/mean x 100).

## Acknowledgments

We thank Laura King, Caroline Nicholas, Renée Kludas, Leanne Redl, Aniruddh Haldar, Felicity Healy, Candice Hall, Elizabeth Crouch, Narelle Shen and Micheala Waibel for their assistance with sample collection and project logistics. We thank Dr Michelle So for critical feedback on the manuscript. We gratefully acknowledge the support of the participants and their families.

## Funding

We thank Operational Infrastructure Support Program of the Victorian Government for support. ML is the recipient of a JDRF Postdoctoral Fellowship (201310741) and a St. Vincent’s Institute Rising Star Award. This work has been supported by: JDRF (1-INO-2022-1124-A-N) and the Medical Research Future Fund (MRFF; RARUR000103).

## Author contributions

Conceptualization: ML, SIM

Methodology: ML, PB, JTD, MYH, SIM

Investigation: ML, PB, LK, AF, MYH, SIM

Visualization: ML, PB, SIM

Funding acquisition: ML, SIM

Resources: JK, JMW, FC, SIM

Supervision: SIM

Writing – original draft: ML, SIM

Writing – review & editing: ML, SIM, JMW, MYH, JTD, FC

## Competing interests

SIM and ML are listed as inventors on provisional patent application 2024903912.

## Data and materials availability

All data are available in the main text or the supplementary materials.

## Notes

### Competing Interest Statement

Authors SIM and ML are listed as inventors in provisional patent application 2024903912.

