## Supplementary Materials for "BASTA, a simple whole blood assay for measuring β-cell antigen specific CD4^+^ T-cell responses in type 1 diabetes"

### Title: A paediatric-friendly whole blood assay detects C-peptide specific CD4$^+$ T-cell responses in Type 1 Diabetes.

**Supplementary Tables**

Supplementary Table 1 Demographics and HLA all blood donors
Supplementary Table 2: Demographics and HLA of pediatric cross-sectional validation cohort
Supplementary Table 3 Sequences of Peptide Stimuli

**Supplementary Figures**

Supplementary Figure 1: CFSE assay and cytokine detection
Supplementary Figure 2: Assay optimization
Supplementary Figure 3: Preliminary assay validation
Supplementary Figure 4: Inter and intra-assay variability
Supplementary Figure 5: Analysis of the IL-2 producing cells
Supplementary Figure 6: Subgroup analysis of IL-2 responses to PPI peptides in pediatric cohort
Supplementary Figure 7: IL-10 production and HLA stratification.

Supplementary Table 1 Demographics and HLA of non-T1D and T1D blood donors.

| | **Non-T1D** | **T1D** |
|---|---|---|
| **No. individual participants** | **44** | **137** |
| Age (years) | 37.23 ± 16.59 (SD)<br>range: 8 - 66 | 13.87 ± 4.25 (SD)<br>range: 3 - 28 |
| Sex (male \| female) | 19 \| 25 | 86 \| 47 |
| Disease duration (years) | N/A | 4.71 ± 4.47 (SD)<br>Range: 0 - 15.72 |
| % HLA-DQ2+ | 11.36 % | 43.07 % |
| % HLA-DQ8+ | 29.55 % | 38.69 % |
| % HLA-DQ2/HLA-DQ8+ | 2.27 % | 18.98 % |
| % Neither HLA-DQ2+ nor HLA-DQ8+ | 54.5 % | 11.7 % |

Supplementary Table 2: Demographics of pediatric participants in cross-sectional validation.

| | **At-risk participants** | **Low-risk pediatric participants** | **New-Onset T1D** | **Established T1D** |
|---|---|---|---|---|
| **No. individual participants** | **21** | **10** | 12 | 20 |
| Age (years) ± (SD) | 6.63 ± 2.3<br>range: 3 – 11 | 15.40 ± 3.89<br>range: 8 - 20 | 11.42 ± 4.23<br>range: 4-17 | 13.35 ± 3.31<br>range: 6 - 17 |
| Sex (male \| female) | 13 \| 8 | 4 \| 6 | 8 \| 4 | 17 \| 3 |
| Disease duration (years) | N/A | N/A | 0.02 ± 0.04 | 7.45 ± 4.60 |
| % HLA-DQ2+ | 38.1 % (8/21) | 0% (0/10) | 58.33% (7/12) | 70% (14/20) |
| % HLA-DQ8+ | 38.1% (8/21) | 20% (2/10) | 41.67% (5/12) | 60% (12/20) |
| % HLA-DQ2/HLA-DQ8+ | 9.5 % (2/21) | 0% (0/10) | 16.67% (2/12) | 35% (7/20) |
| % Neither HLA-DQ2+ nor HLA-DQ8+ | 33.3 % (7/21) | 80% (8/10) | 16.7% (2/12) | 5% (1/20) |
| % ZnT8 AAB+ | 19 % (4/21) | N/A | | N/A |
| % GADA+ | 9.5 % (2/21) | N/A | | N/A |
| % IAA+ | 9.5 % (2/21) | N/A | | N/A |
| % IA-2A+ | 9.5 % (2/21) | N/A | | N/A |
| % Multiple AAB+ | 14.3 % (3/21) | N/A | | N/A |

**Supplementary Table 3: Peptide Sequences**

| Peptide | Sequence | Pool |
|---|---|---|
| **PPI1** | MALWMRLLPLLALLALWG | **POOL1** |
| **PPI2** | LLPLLALLALWGPDPAAA | **POOL1** |
| **PPI3** | LLALWGPDPAAAFVNQHL | **POOL2** |
| **PPI4** | PDPAAAFVNQHLCGSHLV | **POOL2** |
| **PPI5** | FVNQHLCGSHLVEALYLV | **POOL3** |
| **PPI6** | CGSHLVEALYLVCGERGF | **POOL3** |
| **PPI7** | EALYLVCGERGFFYTPKT | **POOL4** |
| **PPI8** | CGERGFFYTPKTRREAED | **POOL4** |
| **PPI9** | FYTPKTRREAEDLQVGQV | **POOL5** |
| **PPI10** | RREAEDLQVGQVELGGGP | **POOL5** |
| **PPI11** | LQVGQVELGGGPGAGSLQ | **POOL6** |
| **PPI12** | ELGGGPGAGSLQPLALEG | **POOL6** |
| **PPI13** | GAGSLQPLALEGSLQKRG | **POOL7** |
| **PPI14** | PLALEGSLQKRGIVEQCC | **POOL7** |
| **PPI15** | SLQKRGIVEQCCTSICSL | **POOL8** |
| **PPI16** | IVEQCCTSICSLYQLEN | **POOL8** |
| **PPI17** | TSICSLYQLENYCN | **POOL8** |
| **Signal Peptide (full-length)** | MALWMRLLPLLALLALWGPDPAAA | **PPI pool** |
| **Insulin B-chain (full-length)** | FVNQHLCGSHLVEALYLVCGERGFFYTPKT | **PPI pool** |
| **C-peptide (full-length 31-mer)** | EAEDLQVGQVELGGGPGAGSLQPLALEGSLQ | **PPI pool** |
| **Insulin A-chain (full-length)** | GIVEQCC*TS*ICSLYQLENYCN | **PPI pool** |

*isoacyl bond for A-chain shown in italic with underline.

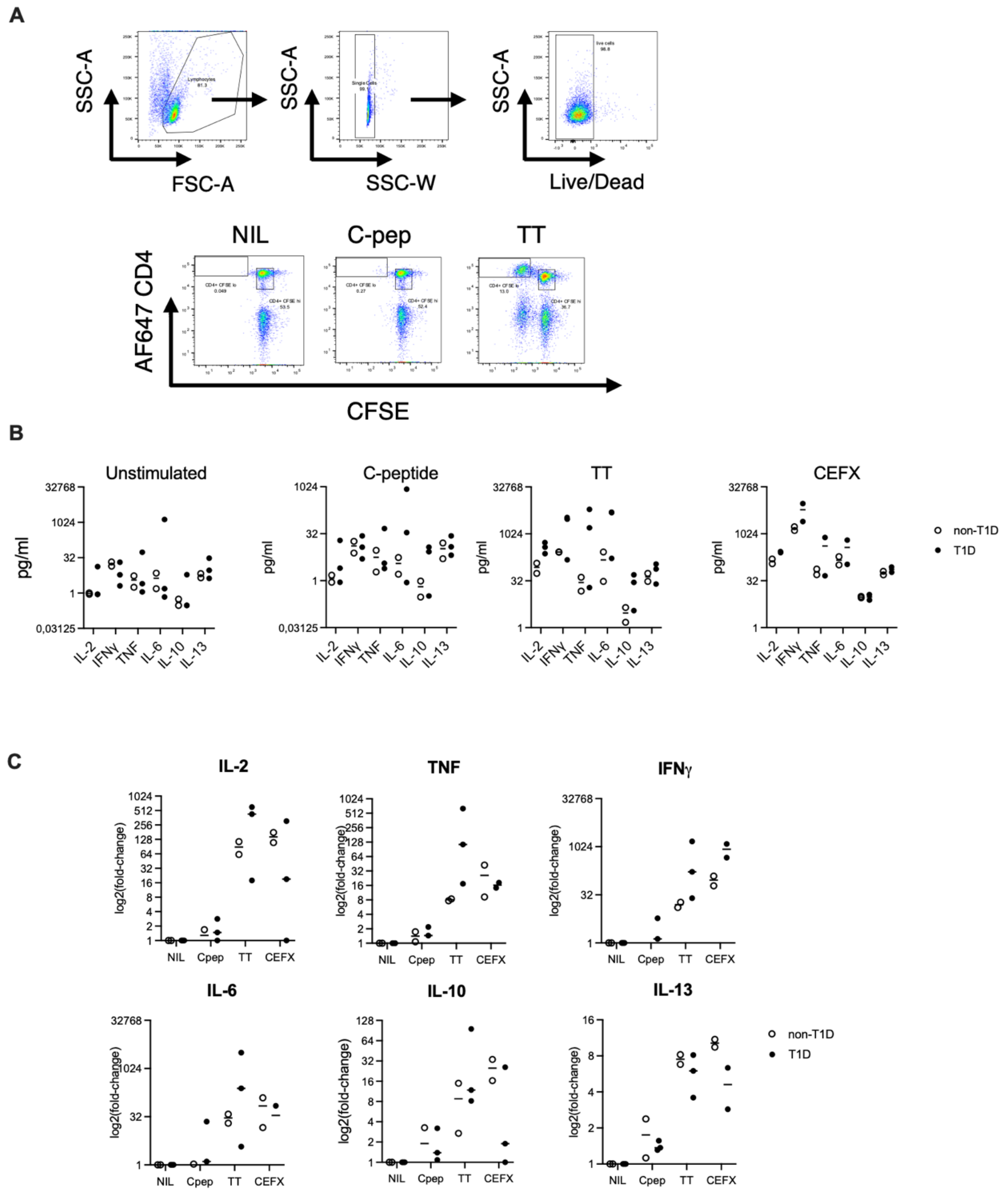


**Fig S1 CFSE assay and cytokine detection (A)** Example FACS plots showing gating and CFSE dye-dilution CD4+ T cell proliferation in untreated versus C-peptide- or TT-stimulated PBMC samples from one subject with newly diagnosed T1D (>3 months). **(B)** Multiplex Cytokine data for whole blood cytokine secretion (IL-2, IFN-gamma, TNF, IL-6, IL-10, IL-13) in blood cultured without treatment, or in response to C-peptide, TT, or pooled positive control peptide epitopes CEFX. **(C)** Cytokine data from (B) represented as a fold-change to levels in unstimulated blood.

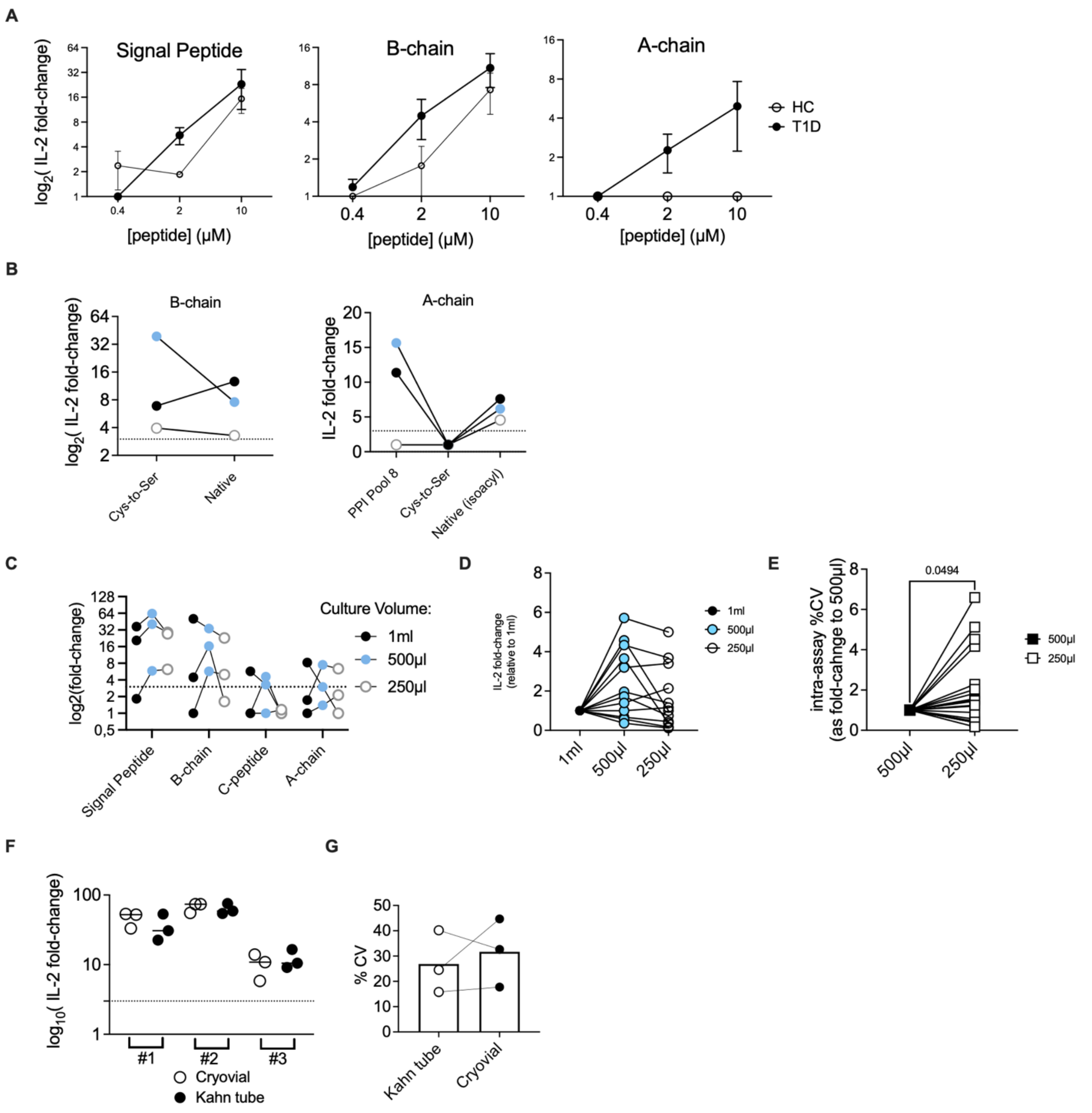


**Fig. S2 Assay optimization** Comparing effect of: (A) Decreasing peptide concentration for full-length PPI Signal peptide, B-chain, and A-chain as stimuli for whole blood IL-2 release. **(B)** Peptide format: Cysteine-to-serine substitutions for A-chain (versus pool of native 14-18mers or full-length native A-chain synthesized as isoacyl form) and B-chain of insulin. **(C)** IL-2 responses from whole blood at 3 culture volumes (either 1ml, 500µl, or 250µl) for each of the four PPI regions for 3 donors with established T1D. **(D)** IL-2 signal strength further compared as relative to 1ml volume for samples in (C). **(E)** Blood cultures performed in triplicate comparing variance (%CV) for 500µl and 250µl culture volumes, using fold-change over %CV at 500µl to compare samples and stimuli. (n = 8 ). **(F)** A combination of all four full-length peptide regions of PPI were tested as a single stimulus, and Cryovials versus 5ml round bottom "Kahn" tubes as culture vessel, with blood from 3 subjects with established T1D assayed in triplicate, and **(G)** %CV per donor compared between vessel type. Dotted line depicts assay cut-off of 3.

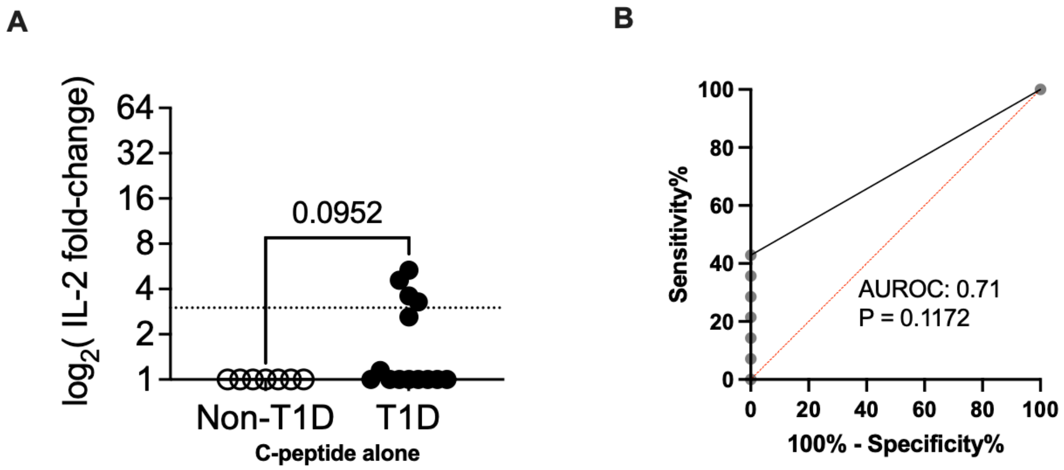


**Fig. S3 Preliminary assay validation (A)** Responses to C-peptide alone in same cohort as 3A, with **(B)** corresponding ROC curve for C-peptide-induced IL-2 values. Dotted line depicts assay cut-off of 3.

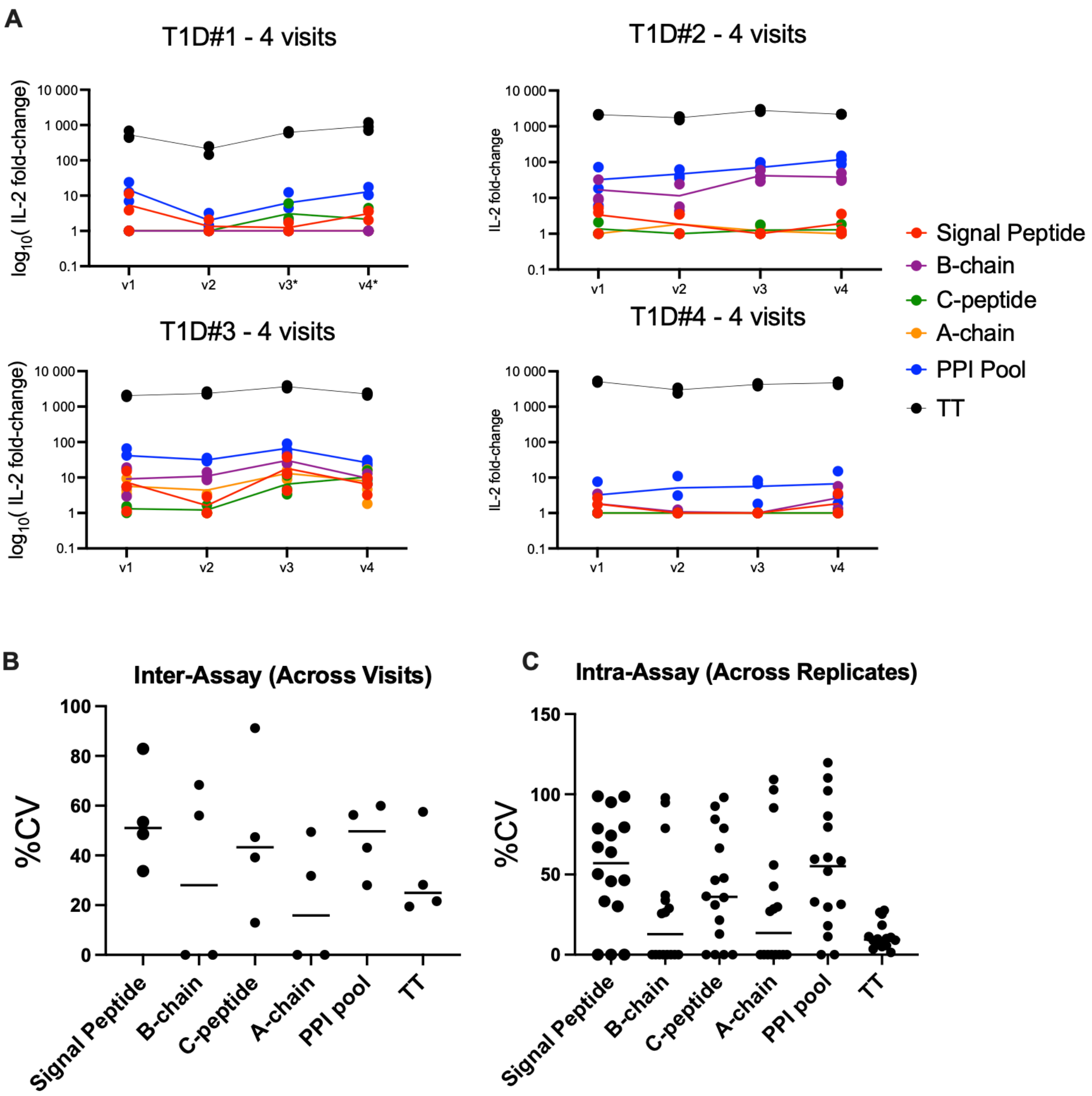


**Fig. S4 Inter and intra-assay variability (A)** Responses to individual full-length peptides from the four PPI regions alongside PPI and TT responses as per (Fig. 4B), measured in triplicate as shown by individual dots, connecting lines show means. **(B)** %CV calculated between triplicates for each stimulus at each visit in (A). **(C)** %CV of mean IL-2 across visits as a measure of inter-assay repeatability for (C), as well as concurrent responses to individual peptides. Lines at median are shown.

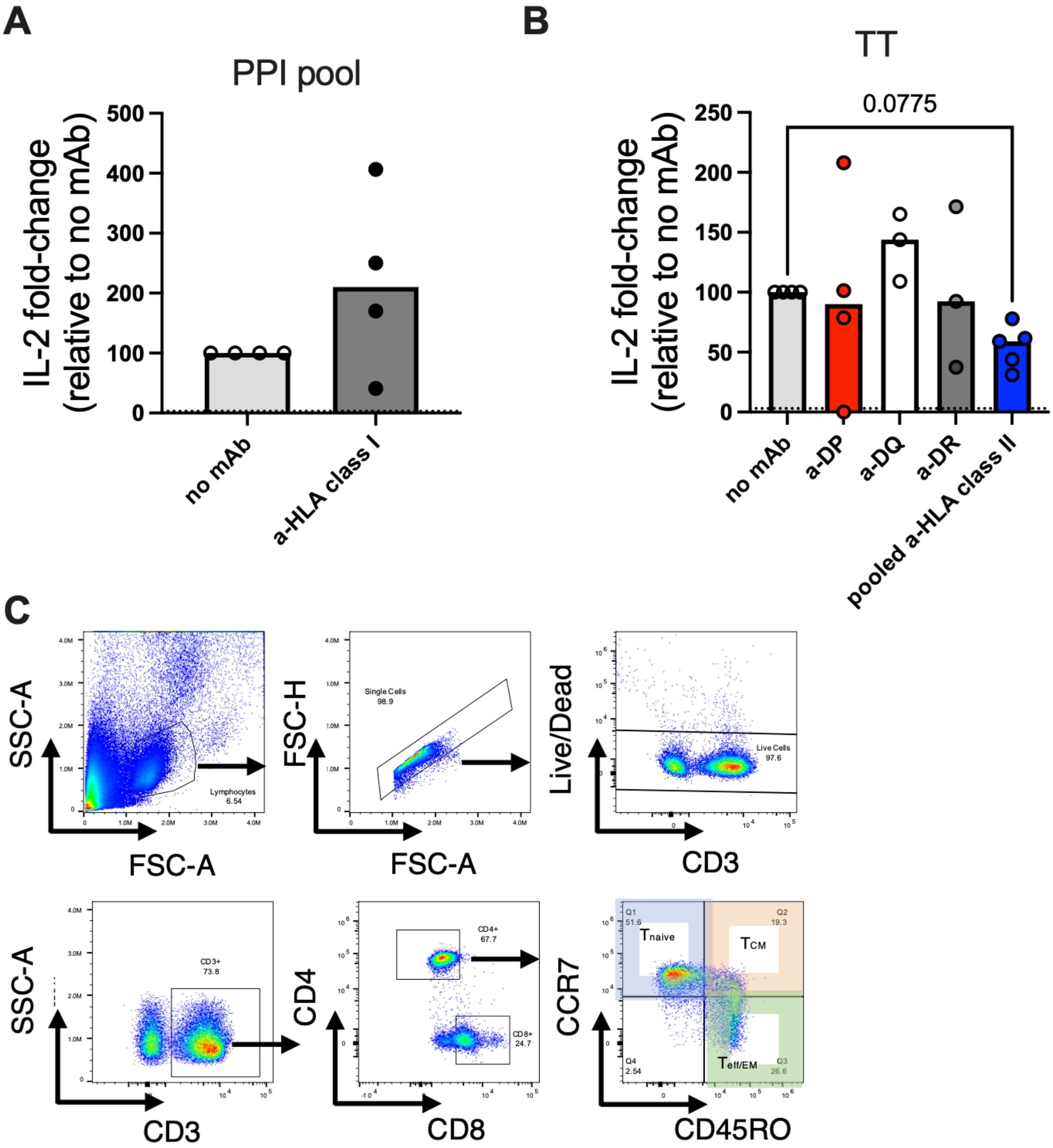


**Fig. S5 Analysis of the IL-2 producing cells (A)** Effect on IL-2 release of blocking antigen presentation via HLA-A, -B, and -C using W6/32 pan -anti-HLA-class-I mAb (2µg/ml) in PPI-stimulated whole blood, relative to PPI-stimulated blood alone. **(B)** mAb inhibition of TT-induced IL-2 as per 4A. **(C)** Representative FACS gating for live CD3+CD4+ T cells and phenotyping using CD45RO and CCR7 for 4B-4D. n= 3-5 per conditions. Bars show medians.

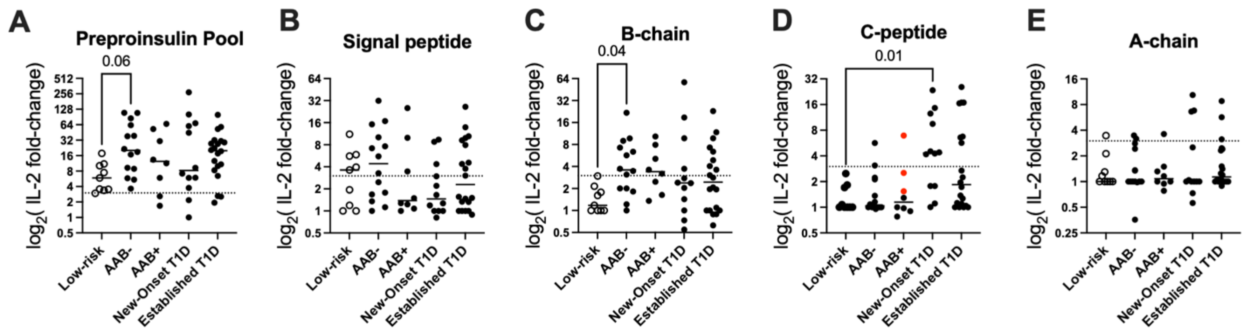


**Fig. S6 Subgroup analysis of IL-2 responses to PPI peptides in pediatric cohort.** IL-2 responses to individual or pooled full-length preproinsulin peptides stratified into five subgroups as described in (Fig. 6A): low-risk, AAB-, AAB+, New-Onset T1D, or established T1D, in response to: **(A)** Preproinsulin pool, **(B)** Signal peptide, **(C)** B-chain, **(D)** C-peptide (red circles = multi-AAB+ individuals), **(E)** A-chain. Dotted line represents 3-fold over background. Lines at median are shown.

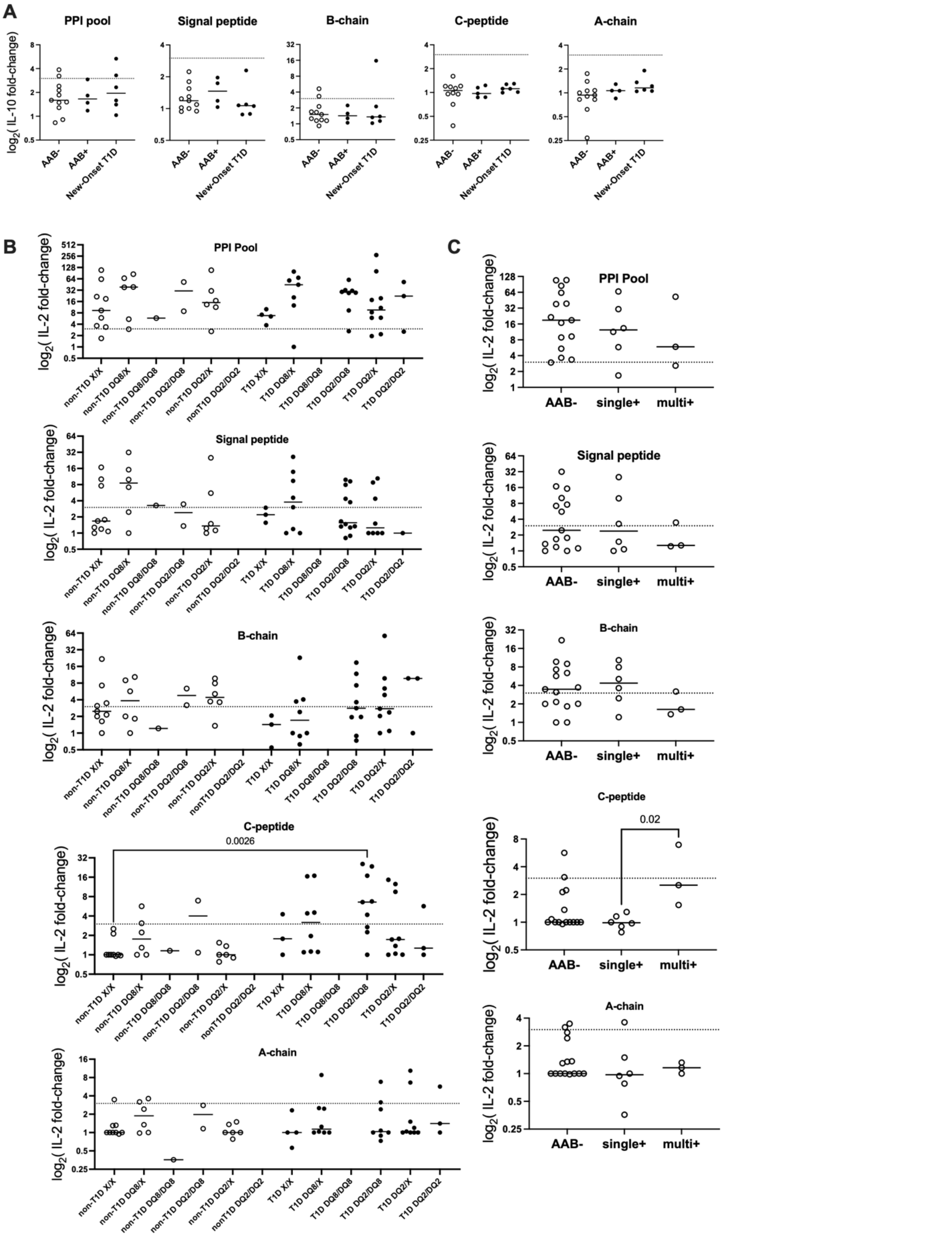


**Fig. S7 IL-10 production, HLA and AAB stratification (A)** IL-10 responses from AAB- and AAB+ at-risk paediatric donors from 5A, and subset of participants with New-Onset T1D. **(B)** Stratification of IL-2 responses in 5C according to HLA-type relative to high-risk HLA-class II allelic haplotypes DQ2, DQ8, with non-T1D associated variants represented as X. **(C)** Stratification of IL-2 responses in 5C in at-risk ENDIA subjects according to AAB-, single AAB+ (single+) or multi-AAB+ (multi+) antibody status. Dotted line represents 3-fold over background. Lines at median are shown.
